# CCL3 promotes germinal center B cells sampling by follicular regulatory T cells

**DOI:** 10.1101/295980

**Authors:** Zachary L. Benet, Matangi Marthi, Rita Wu, Jackson S. Turner, Jahan B. Gabayre, Michael I. Ivanitskiy, Sahil S. Sethi, Irina L. Grigorova

**Affiliations:** Department of Microbiology and Immunology, University of Michigan Medical School, 1150 West Medical Center Drive, 5641 Medical Science II, Ann Arbor, Michigan 48109-5620

**Author notes:** Please address correspondence to: Irina L. Grigorova, 1150 W. Medical Center Dr, 6748 MSII Ann Arbor, MI 48109, (p), (f), ORCID: 0000-0002-4963-7403.

## Abstract

Previous studies and our findings suggest upregulated expression of proinflammatory chemokines CCL3/4 in germinal center (GC) centrocytes. However, the role of CCL3/4 for centrocyte interactions with follicular T cells and regulation of humoral immunity is poorly understood. We found that CCL3 promotes chemotaxis of Tfr cells *ex vivo. In vivo* CCL3 is not required for Tfr cells recruitment of into the GC light zone. However, B cells-intrinsic production of CCL3 promotes their direct interactions and negative regulation by follicular regulatory T cells (Tfr) within GCs.

## INTRODUCTION

CCL3 and CCL4 (MIP1-α and MIP1-β) are proinflammatory chemokines that are secreted by various types of immune cells upon activation and play important roles in inflammatory responses and multiple other processes (Menten, Wuyts et al. 2002, Maurer and von Stebut 2004). In B cell cultures with naive, memory, or GC B cells, the cross-linking of B cell receptors (BCRs) upregulates expression and secretion of CCL3/4 (Krzysiek, Lefevre et al. 1999, Bystry, Aluvihare et al. 2001). In addition, analysis of published GC microarray data suggests that expression of CCL3/4 may be elevated in GC centrocytes compared to centroblasts (Caron, Le Gallou et al. 2009, Compagno, Lim et al. 2009, Victora, Dominguez-Sola et al. 2012). Despite multiple indications that CCL3/4 is secreted by activated and GC B cells, the significance of B-cell intrinsic production of proinflammatory chemokines for regulation of humoral response is unclear.

In 2001 Bystry et al. demonstrated that a subset of splenic CD4 T cells that were CD25^high^, expressed high levels of TGF-β and CTLA-4, and had a suppressive phenotype *ex vivo*, could migrate to CCL4 (but not CCL3) in transwell assays. The above observations raised the possibility that CCL4 may promote Tregs’ interactions with activated B cells or dendritic cells (which also produce CCL3/4) to regulate B cell responses. However, whether more recently identified subset of follicular resident Tregs can respond to CCL3 or CCL4 has been unclear (Lim, Hillsamer et al. 2005, Alexander, Tygrett et al. 2011, Chung, Tanaka et al. 2011, Linterman, Pierson et al. 2011).

Tfr cells are a subset of FoxP3^pos^ Tregs that play a role in the control of GC responses. While the majority of Tfr cells arise from natural Tregs, some can be induced from foreign antigen-specific Th cells (Aloulou, Carr et al. 2016). Similarly to Tfh cells, Tfr cells develop in the secondary lymphoid organs following foreign antigen challenge, express the transcription factor Bcl6, upregulate surface expression of CXCR5, PD1 and ICOS receptors, and localize to the follicles and the GCs (Chung, Tanaka et al. 2011, Linterman, Pierson et al. 2011, Wollenberg, Agua-Doce et al. 2011). Deficiency in Tfr cells has been reported to induce a 1.5 to 2-fold increase in GC size at the peak of GC response (Chung, Tanaka et al. 2011, Linterman, Pierson et al. 2011, Wollenberg, Agua-Doce et al. 2011), to affect Tfh cell’s cytokine production and Ab class-switching (Sage, Francisco et al. 2013, Sage, Paterson et al. 2014, Wu, Chen et al. 2016), to increase recruitment of non-foreign antigen (Ag)-specific B cell clones into GCs (Linterman, Pierson et al. 2011, Wu, Chen et al. 2016), and to promote development of self-reactive antibodies (Wu, Chen et al. 2016, Botta, Fuller et al. 2017) and even autoimmunity in influenza-infected mice (Fu, Liu et al. 2018). While multiple mechanisms of Tfr cells’ action have been suggested based on the *in vivo* and *ex vivo* studies (Alexander, Tygrett et al. 2011, Sage, Francisco et al. 2013, Sage, Paterson et al. 2014, Wing, Ise et al. 2014, Wing and Sakaguchi 2014, Sage, Ron-Harel et al. 2016), whether *in vivo* Tfr cells only regulate Tfh cells and thus indirectly control GC responses or can also act on the GC B cells directly remains an open-ended question.

Here we investigated the role of B cell-intrinsic production of CCL3 in the regulation of GC responses. Based on 2-photon imaging of murine lymph nodes we found that production of CCL3 by foreign Ag-specific GC B cells promotes their sampling and direct inhibition by Tfr at the peak of GC response.

## RESULTS

### A small subset of GC centrocytes upregulates expression of CCL3 and CCL4

Based on the previous microarray data (Caron, Le Gallou et al. 2009, Compagno, Lim et al. 2009, Victora, Dominguez-Sola et al. 2012) GC centrocytes (CC) may have elevated expression of CCL3/4 compared to centroblasts (CB). To verify expression of *Ccl3* and *Ccl4* in murine GC B cells, we performed qRT-PCR analysis of GC CC, CB and non-GC B cells sorted from the draining lymph nodes (dLNs) of immunized mice at 10 days post immunization (d.p.i) (**Fig. 1a**). Consistent with previous microarray analysis, we found that expression of CCL3/4 is upregulated in murine GC CC compared to CB (**Fig. 1b, c**).

**Figure 1.**
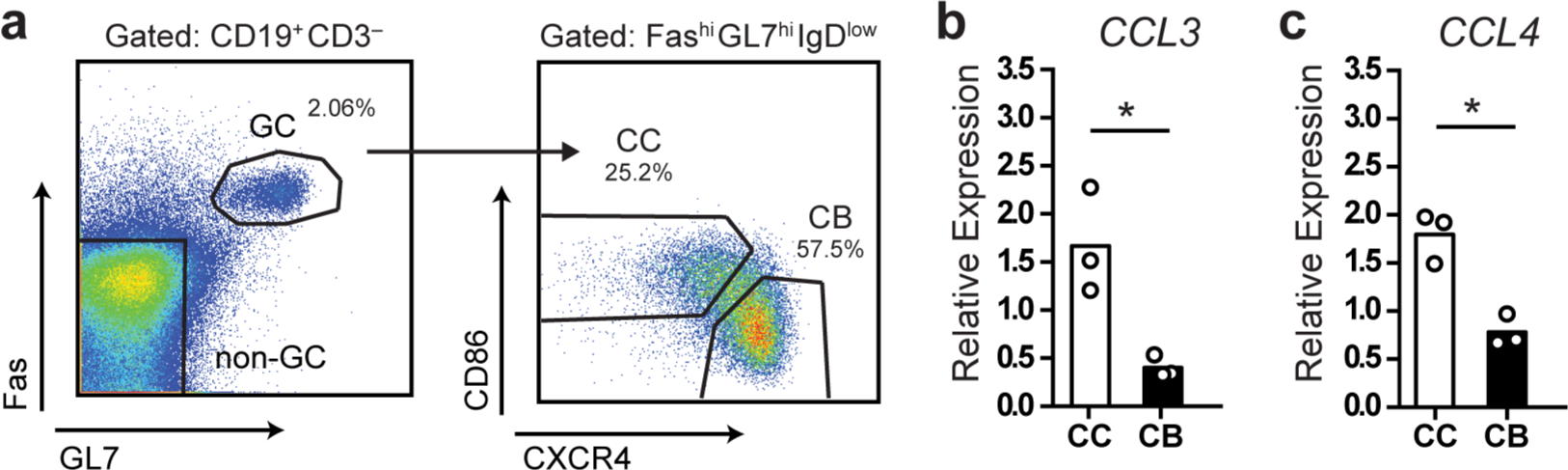
A small subset of GC centrocytes upregulates CCL3/4 expression. qRT-PCR analysis of CCL3 and CCL4 expression in GC and non-GC B cells sorted from the draining LNs of mice s.c. immunized with 50 *μ*g ovalbumin (OVA) in Ribi at 10 days post immunization (d.p.i). **a**, The gating strategy for cell sorting. **b, c**, Relative expression of CCL3/4 in bulk GC centroblasts (CB) and centrocytes (CC) as normalized to non-GC B cells (FAS^low^ GL7^low^). Chemokine C_T_ values were normalized to β2m C_T_. Bars represent mean with dots representing average values per experiment. Data represent n=3 independent experiments, 4 mice total. *, P<0.05, two-tailed Student’s *t*-test.

### CCL3 and CCL4 induce chemotaxis of follicular regulatory T cells *ex vivo*

To determine whether CCL3/4 may promote chemotaxis of follicular T cells *ex vivo* we performed transwell migration analysis of CD4 T cells isolated from dLNs of mice at 10 d.p.i. with OVA in Ribi adjuvant. CXCR5^high^ PD1^high^ FoxP3+ Tfr cells transmigrated in response to CCL3 chemokine (**Fig. 2a, b**). Similar trends were observed for CXCR5^low^ PD1^low^ FoxP3^+^ and CXCR5^int^ PD1^int^ FoxP3+ CD4 T cells (**Fig. 2a, b**). We also observed transmigration of Tfr and other regulatory T cell subsets to CCL4 (**Fig. 2c**). The observed transmigration was predominantly due to chemotaxis rather than chemokinesis, since addition of CCL3 or CCL4 chemokines to both the upper and the lower wells of the transwell chamber did not promote Tregs’ transmigration (**Fig. 2b, c**). No significant transmigration of Tfh cells (CXCR5^high^ PD1^high^ FoxP3^−^) to CCL3 and CCL4 chemokines was observed (**Fig. 2d, e**). These data suggest that CCL3 and CCL4 can induce chemotaxis of Tfr and possibly other Treg cells *ex vivo.*

**Figure 2.**
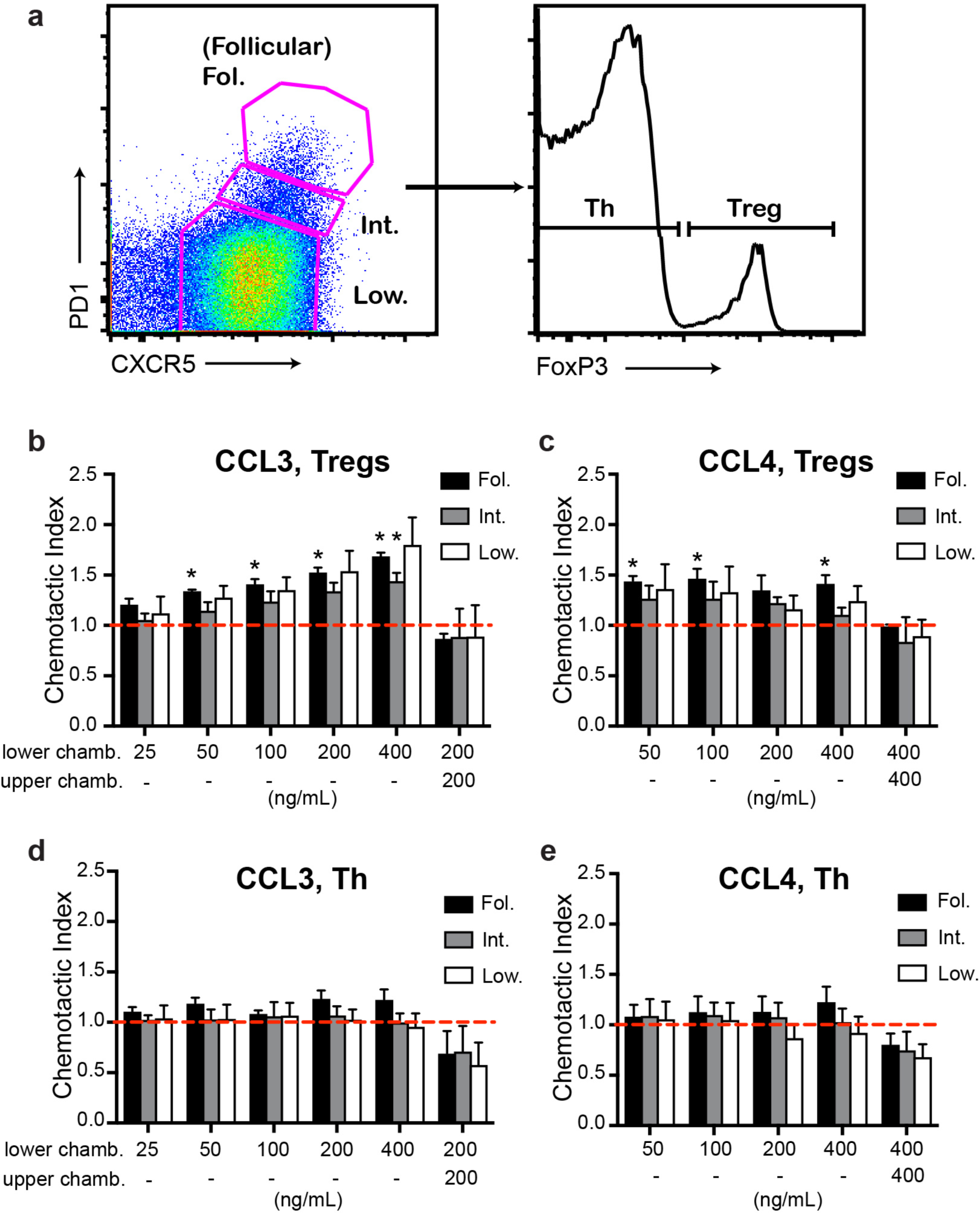
Chemotaxis of Tfr cells to CCL3 and CCL4 *ex vivo.* a-e, *Ex vivo* transwell assay of CD4^+^ T cells purified from dLNs of WT mice at 10 d.p.i. with 100 *μ*g OVA in Ribi and analyzed by flow cytometry using the gating strategy as in (a). Transmigration of CXCR5^high^PD1^high^ (Fol., black bars), CXCR5^int^PD^int^ (Int., grey bars), or CXCR5^low^PD1^low^ (Low, white bars) CD4^pos^ CD8^neg^ B220 ^neg^ cell populations that also express Foxp3 (b, c) or were FoxP3 ^neg^ (d, e) were measured against CCL3 (b, d) and CCL4 (c, e). The chemokines were added either to the lower chambers (chamb.) of transwells for analysis of chemotaxis or to both the upper and the lower chambers for analysis of chemokinesis. Chemotactic index was calculated as the ratio of cells that transmigrated towards the chemokine vs. no chemokine control (dashed red line). Chemotaxis and chemokinesis data are derived from 3 and 2 independent experiments correspondingly with 2 mice per experiment. Bars represent mean ± SEM, *, P<0.05, two-tailed, one-sample Student’s *t*-test (compared to 1).

### CCL3 does not recruit Tfr cells into the GC light zone

To determine if CCL3/4 play a role in the regulation of GC size, we utilized CCL3-KO mice (Cook, Beck et al. 1995). In unimmunized CCL3-KO and WT mice we observed no significant difference in the GC B cells numbers in peripheral lymph nodes (pLNs), spleens, mesenteric LNs (mLNs) and peyer patches (PP) (**Fig. 3a**). However, at 10 d.p.i. we detected a small, but significant increase in the GC response in the dLNs of CCL3-KO mice compared to WT that was independent of the antigen or the adjuvant used (**Fig. 3b-e**). We then tested whether in littermate-control CCL3^−/−^ mice GCs were also elevated compared to CCL3^+/+^, and confirmed the observed phenotype (**Fig. 3f, g)**. Interestingly, treatment of WT or CCL3-KO mice with CCL4-neutralizing antibodies during formation of GCs, at 7 d.p.i., did not lead to further increase in the GC response (**Supplementary Fig. 1**). The observed accumulation of GC B cells at 10 d.p.i. was not due to significant changes in formation of Tfh and Tfr cells or in their ratio (**Fig. 3h-j**). In addition, the increase in GC B cells did not lead to significant rise in the numbers of plasmablasts (PB) at 10 d.p.i. (**Fig. 3k, l**).

**Figure 3.**
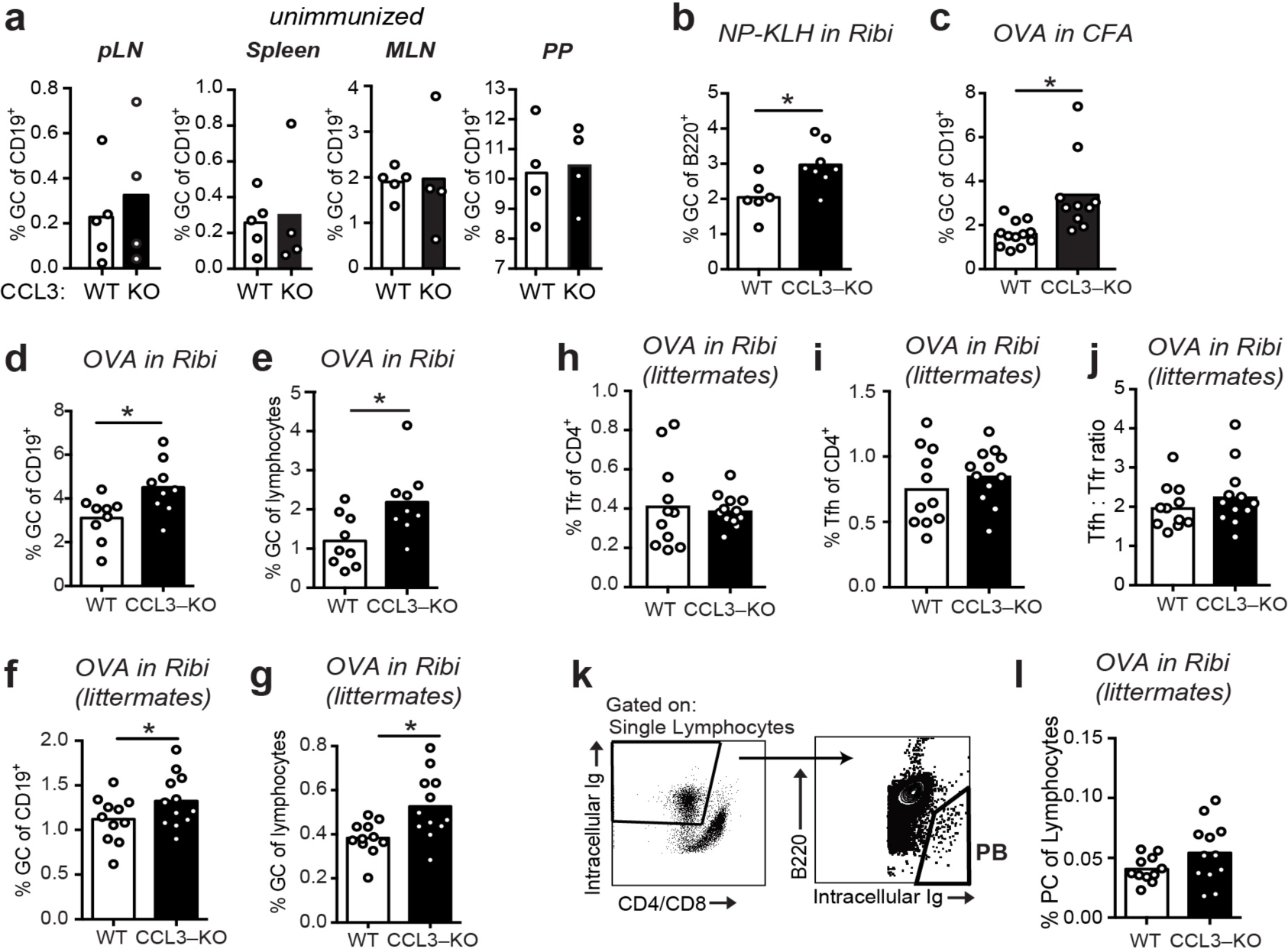
Increased GC response in CCL3-K0 mice following immunization. **a-g**, Flow cytometry analysis of the GC B cells (Fas^high^ GL7^high^ lgD^l0W^) as a fraction of total B cells (a-d, f) or total lymphocytes (e, g) in WT and CCL3-KO mice, **a**, Analysis of GCs in pLNs (inguinal, brachial and auxiliary LNs combined), spleens, MLNs, and PP from unimmunized mice, **b-g**, Analysis of GCs in the dLNs from mice s.c. immunized with 50 /vg of 4-Hydroxy-3-nitrophenyl (*NP)* acetyl-*hapten* conjugated to *Keyhole limpet hemocyanin* (KLH) in Ribi adjuvant (b) or 50 /vg of OVA in Complete Fruend’s Adjuvant (CFA) (c) or 50 /vg OVA in Ribi (d-g) at 10 d.p.i. **f-l**, Flow cytometry analysis of GC (f, g), Follicular T cell (h-j), and plasma cell (PC) (k, I) response in dLNs of WT and CCL3-KO littermate conrol mice s.c. immunized with 50 /vg OVA in Ribi at 10 d.p.i. **h-j**, Tfr (CXCR5^high^ PD1^high^ FoxP3^pos^ in h) and Tfh (CXCR5^high^ PD1^high^ FoxP3^neg^ in i) cells as a fraction of CD4 T cells, **k, I**, PB gating strategy (k) and PBs as a fraction of total lymphocytes (I). Each symbol represents one mouse. Bars represent mean. Data are derived from 2 or 3 independent experiments. *, P<0.05, Student’s *t*-test (two-sided in b-e, one-sided in f, g).

Our observations that CCL3 promotes Tfr cells chemotaxis *ex vivo* and plays a role in the control over GC size *in vivo* led us to ask whether CCL3 produced by GC centrocytes recruits Tfr cells from the follicles into the GCs. If this hypothesis is correct, the frequency of Tfr cells in the GC light zone should be reduced in CCL3-KO mice compared to WT mice. To test this, we analyzed the density of Tregs in the GCs (both in the light and the dark zones) and in the follicles of the draining LNs from immunized CCL3-KO and WT mice. Fixed LNs were sectioned and stained with fluorescently conjugated antibodies towards IgD, CD35, CD4, and FoxP3 and analyzed by confocal microscopy (**Figure 4a-c**). In WT mice, CD4^+^ FoxP3^+^ Treg cells were enriched in the follicles compared to the GCs (**Figure 4c, d**). In contrast to the expected decrease in the Tfr cell frequency within the GC light zone of CCL3-KO mice, the density and recruitment index of CD4^+^ FoxP3^+^ cells in the light zones of CCL3 KO and WT mice were comparable (**Figure 4d, e**). However, the recruitment index calculated for Tregs’ access into the GC dark zone relative to the follicle was higher for CCL3-KO mice (**Figure 4e**). Interestingly, we also observed modest enrichment of Tfh cells in the dark zones of GCs in CCL3-KO mice (**Figure 4f, e**). Based on this data we conclude that CCL3/4 is not required for Tfr cells’ recruitment into the GC light zone from the follicles, but may play a role in limiting access of follicular T cells to the GC dark zone.

**Figure 4.**
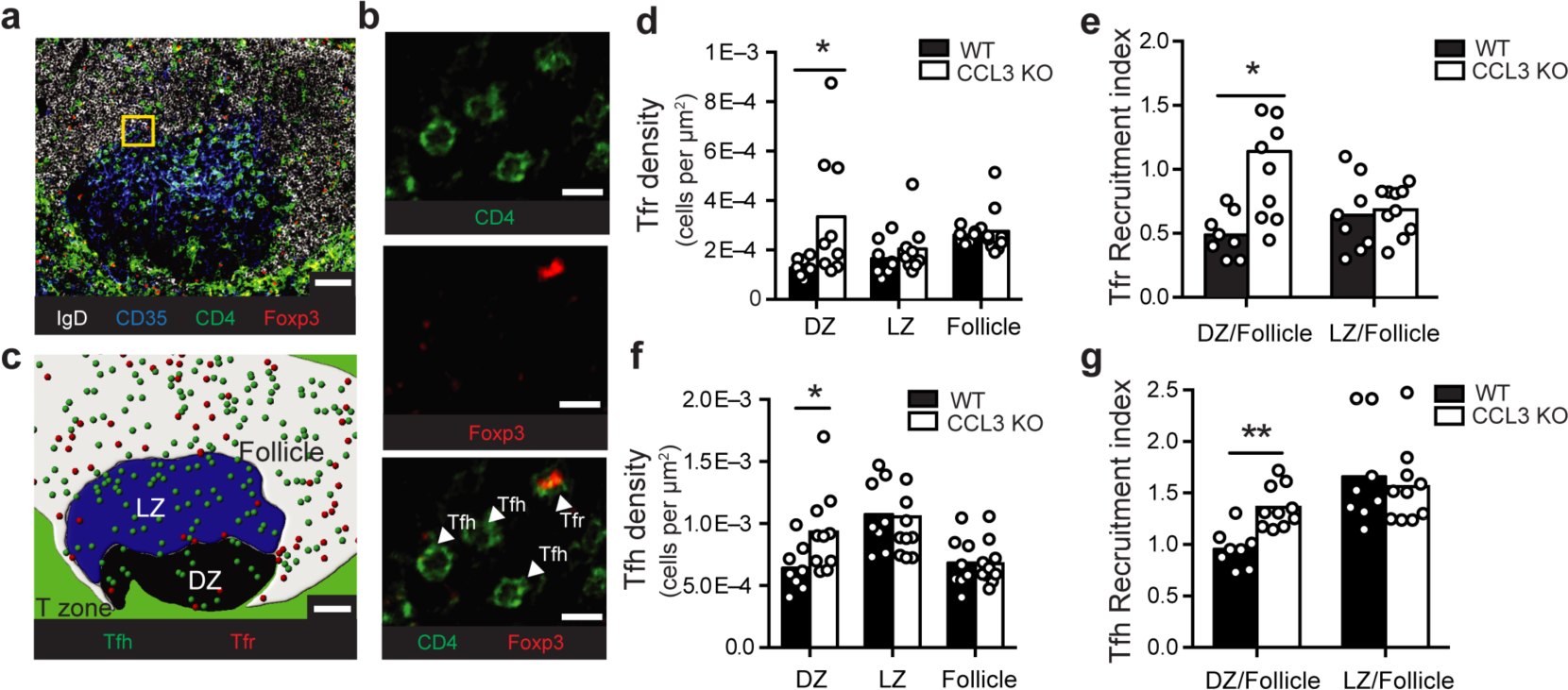
CCL3 does not recruit Tfr or Tfh cells into the GC light zone. Immunofluorescent analysis of Tfr and Tfh cell localization in the GCs and follicles of draining pLNs from WT or CCL3-KO mice at 10 d.p.i. with 50 μg OVA in Ribi. **a-c**, Representative example of a confocal image of pLN section and its analysis, **a**, Confocal image of a GC-containing 10 μm section of pLN from WT mouse, stained with antibodies against IgD (white), CD35 (blue), CD4 (green) and Foxp3 (red), **b**, Magnified image from the inset (in **a)** that illustrates how Treg (CD4^+^Foxp3^+^) and Th (CD4^+^Foxp3^−^) cells were identified, **c**, Reconstruction of GC light zone (blue area), dark zone (black area), follicle (white area), Tregs (red circles) and Th cells (green circles) within the confocal image shown in **(a)** using manually defined surfaces in Imaris. Scale bars - 50 μm in **a, c** and 10 μm in **b. d-g** Quantitative analysis of Treg **(d, e)** and Th **(f, g)** cell abundance in the GCs and the follicles. Density of Tregs **(d)** and Th cells **(f)** in GC dark zone (DZ), light zone (LZ) and follicles around GCs calculated as the number of T cells in each zone normalized to the total area of that zone. Recruitment index calculated as the density of Tregs **(e)** or Th cells **(g)** in GC DZ or LZ normalized to their density in the GC-containing follicle, **d-g**, Each symbol represents analysis of pLN section with a distinct GC. Data represents n=3 independent experiments with 4 total mice per genotype. Bars represent mean. *, P<0.05, two-tailed Student’s *t*-test.

### Adoptively transferred Tregs can become Tfrs’ and be visualized by 2P microscopy

We then asked whether CCL3 could promote individual interactions of Tfr cells with GC B cells *in vivo.* To directly test this, we developed an imaging strategy that enabled visualization of adoptively transferred Tregs within the GC-containing follicles of living mice using 2-photon (2P) microscopy (**Fig. 5a**). Because intravenous injection of Tregs is insufficient to get enough of the transferred Tregs into peripheral LNs for microscopy analysis, we utilized the fact that adoptively transferred Tregs undergo proliferation in recipient FoxP3^DTR^ mice upon diphtheria toxin (DTx)-induced ablation of DTx receptor (DTR)-expressing resident Tregs (Kim, Rasmussen et al. 2007). Of note, while treatment of FoxP3^DTR^ mice with DTx leads to development of severe autoimmune disease, pre-transfer of 10^6^ polyclonal Tregs rescues FoxP3^DTR^ mice from autoimmunity (Kim, Rasmussen et al. 2007). To generate mice with a high number of brightly fluorescent Tfr cells, we first transferred 10^6^ polyclonal Tregs that expressed both Foxp3-GFP and tdTomato into Foxp3^DTR^ mice (Bettelli, Carrier et al. 2006, Kim, Rasmussen et al. 2007). We then treated the Foxp3^DTR^ recipients with 5 μg of DTx two times, one week apart. As reported before, following transient ablation of Foxp3^DTR^ Tregs, the adoptively transferred Tregs as well as the remaining endogenous Tregs underwent vigorous proliferation. At 14 days following the initial DTx treatment, tdTomato Tregs represented about 50% of all Tregs in the blood (**Fig. 5b**). We then co-transferred Duck Egg Lysozyme (DEL)-specific HyHEL10 B cells expressing CFP, OVA-specific OTII CD4 T cells expressing GFP, as well as non fluorescent HyHEL10 B and OTII T cells into the same recipient mice and induced their recruitment into the GCs by s.c. immunization with DEL-OVA as previously described (Allen, Okada et al. 2007). By 8 d.p.i. the overall levels of Tregs in the blood had returned to normal (**Fig. 5b**). Based on confocal and flow cytometry analysis we determined that over a fifth of CD4^+^Foxp3^+^ cells from dLNs expressed tdTomato and that tdTomato^+^ cells were almost exclusively Foxp3^+^ (**Fig. 5c, d**). Additionally, tdTomato^+^ Foxp3^+^ cells that were also CXCR5^high^ PD1^high^ had increased expression of Bcl6^+^ as expected for Tfr cells (**Fig. 5c**). At 7-8 d.p.i. a relatively minor fraction of GC-proximal Tfr cells entered into the GCs, while majority of the cells moved around GCs in proximity to the outer edge GC B cells (**Fig. 5e, Supplementary Movie 1**). These data suggest that the adoptive transfer of fluorescent Tregs followed by transient ablation of non-fluorescent endogenous Tregs and immunization is sufficient to visualize Tfr cells within GC-associated follicles in living mice by 2P microscopy.

**Figure 5.**
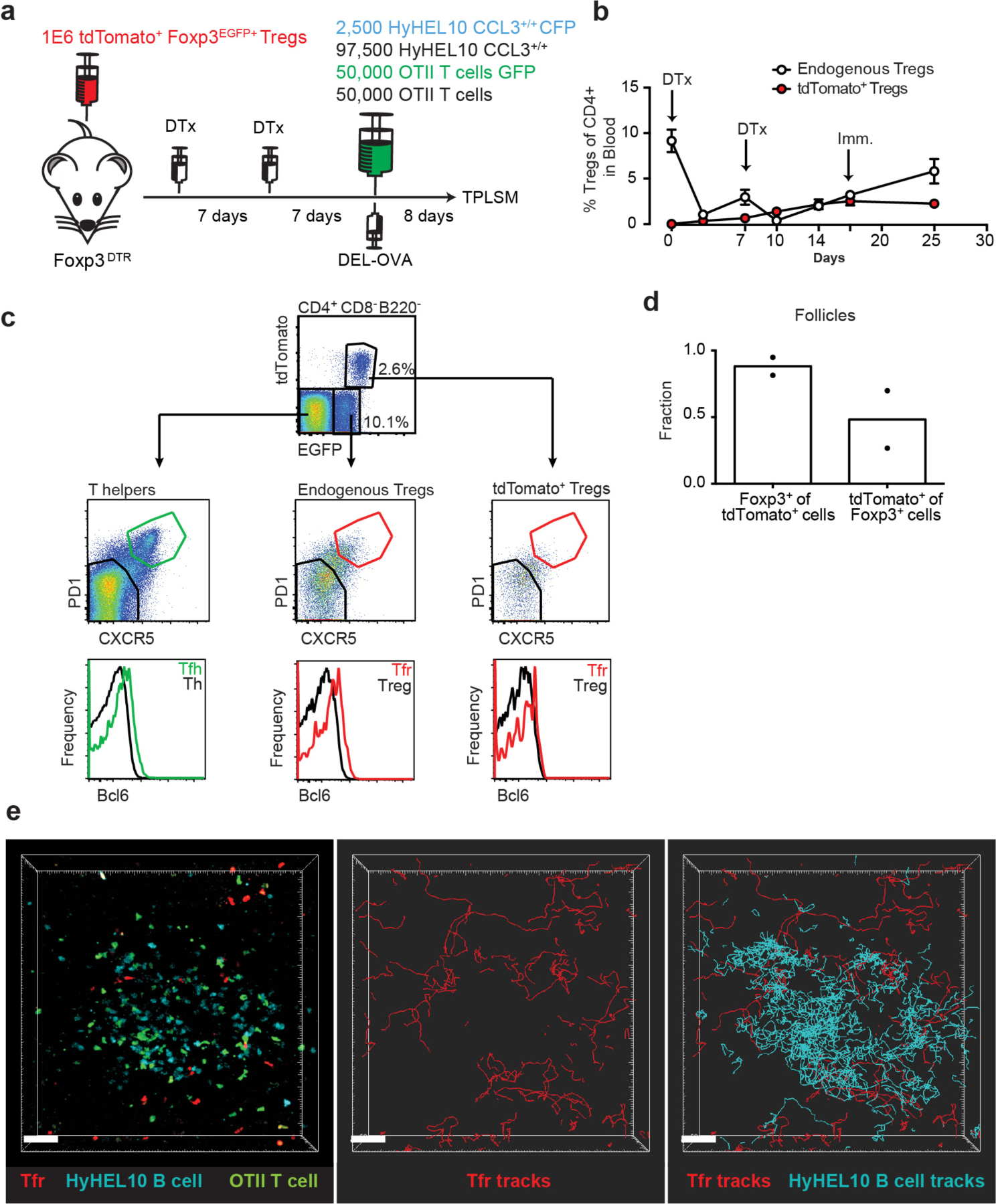
2P visualization of Tfr cells and HyHEL10 GC B cells. a, Experimental diagram for generation of recipient mice with large numbers of highly fluorescent tdTomato^+^ Tregs and visualization of Tfr and HyHEL10 (Hy10) GC B cells in the dLNs by 2P microscopy. **b**, Flow cytometry analysis of the endogenous and transferred TdTomato^+^ Treg numbers in the blood of FoxP3^DTR^ mice throughout their treatments with DTx and immunization, performed as in (a). **c-e**, Analysis of dLNs from mice generated through the procedure described in (a) at 8 d.p.i. **c**, Representative flow cytometry analysis of Bcl6 expression in the CXCR5^low^PD1^low^ and CXCR5^high^PD1^high^ subsets of tdTomato^+^ Tregs (top panels), endogenous Tregs (middle panels) and endogenous Th cells (bottom panels). **d**, Confocal immunofluorescence analysis of dLN sections for the fraction of tdTomato^+^ cells that express Foxp3 (left bars) and fraction of Foxp3^+^ cells that express tdTomato^+^ (right bars) in GC-containing follicles.**e**, Representative 2P imaging analysis of Tfr cells localization with respect to GCs. A snapshot (left panel) from an intravital imaging experiment as analyzed in Imaris software. Cell trajectory analysis for Tfr cells (middle panel) and both Tfr cells and GC B cells (right panel). Scale bars 40 μιτι. Data are from 2 independent experiments.

### GC B cells’ CCL3 promotes their contacts with Tfr, but not Tfh cells *in vivo*

To determine whether GC B cells’ intrinsic production of CCL3 promotes their direct interactions with Tfr cells or with Tfh cells *in vivo* we utilized the experimental setup developed by us (**Fig. 5**) and in previous work (Allen, Okada et al. 2007) and outlined in **Figure. 6a** and **b**. For analysis of T cell interactions with CCL3^+/+^ and CCL3^−/−^ GC B cells, recipient mice were co-transferred with CCL3^+/+^ HyHEL10 B cells that expressed CFP and CCL3^−/−^ HyHEL10 B cells that expressed GFP. To determine Tfr cells’ interactions with GC B cells we tracked tdTomato+ Tfr cells when they contacted or passed through the GCs, and analyzed their interactions with fluorescent CCL3^+/+^ and CCL3^−/−^ foreign Ag-specific HyHEL10 B cells within the same GCs (**Fig. 6c**,

**Figure 6.**
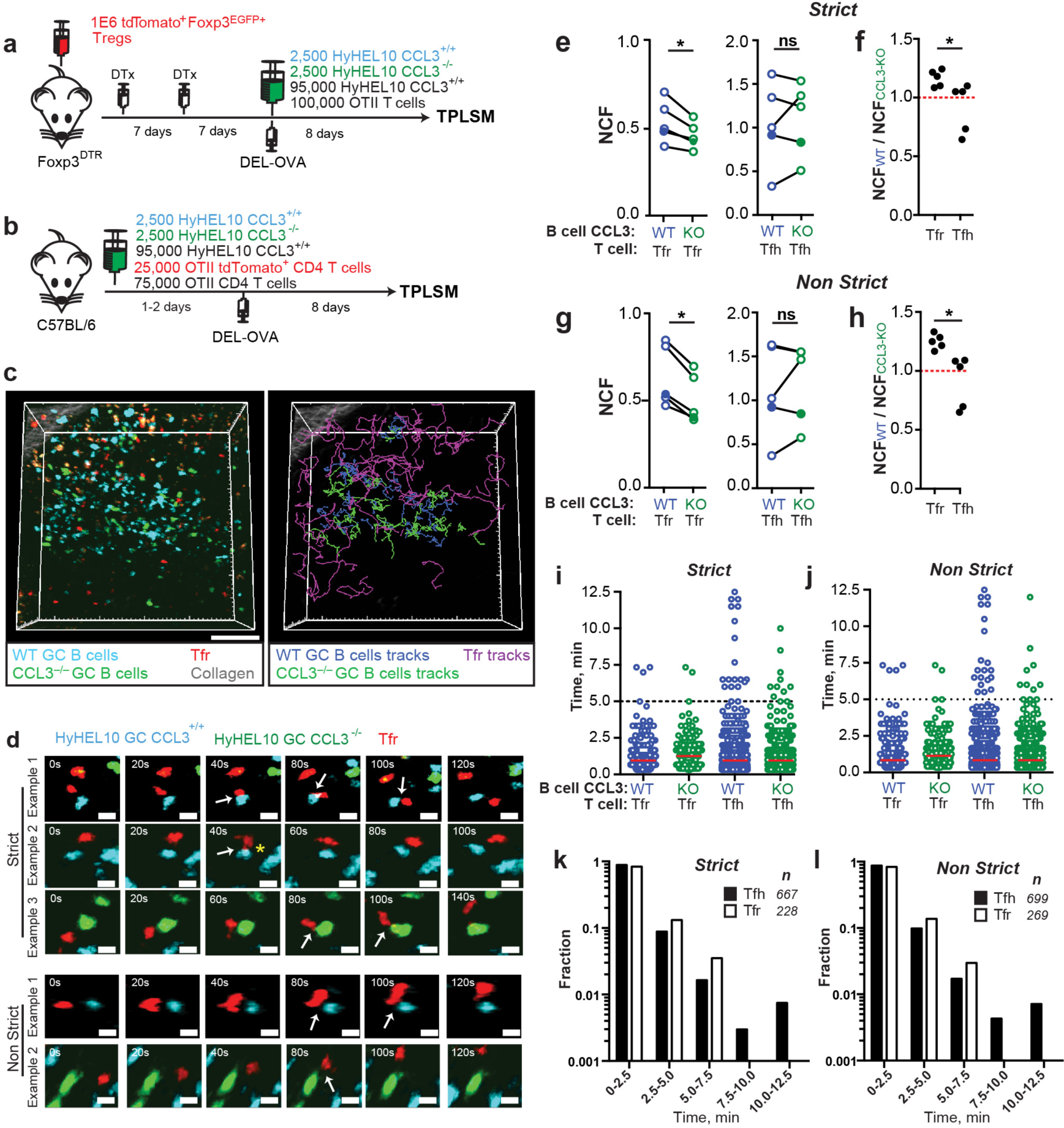
Tfr cells make less frequent contacts with CCL3-KO B cells in GCs. 2P imaging analysis of Tfr and Tfh cells contacts with CCL3^+/+^ CFP and CCL3^−/−^ GFP HyHELI 0 GC B cells in the dLNs of mice at 8 d.p.i. **a, b**, Experimental diagrams for imaging GC B cell interactions with Tfr (a) and Tfh (b). **c**, A snapshot (**left panel**) and cell trajectory analysis (**right panel**) from an intravital imaging experiment performed as described in a. Scale bars - 50 μm. Auto-fluorescent cells are orange. **d**, Time-lapse images of Tfr cells interacting with CCL3^+/+^ or CCL3^−/−^ HyHEL10 B cells within GCs. Cell contacts were verified in 3D space, classified as Strict (definitive contacts) and Non Strict (all possible contacts) and indicated by white arrows. Yellow star illustrates pseudopod extension by a Tfr cell towards CFP GC B cell. Images are displayed as 20 *μ*m z-stacks. Scale bars - 5 μm. **e-j**, Quantitative analysis of CCL3^+/+^ (blue circles) and CCL3^−/−^ (green circles) HyHEL10 GC B cell interactions with Tfr or Tfh cells. Closed symbols represent intravital and opened symbols - explanted dLNs imaging. The data was analyzed in a blinded fashion to avoid possible bias in cell contact definition. **e, g**, Normalized contact frequency (NCF) calculated for Strict (e) and Non Strict (g) as the number of Tfr or Tfh cells’ interactions with CCL3^+/+^ and CCL3^−/−^ HyHEL10 B cells within defined volume of GCs normalized to the average number of HyHEL10 cells of each genotype. Linked symbols correspond to GC B cells in the same movie. *, P<0.05, Wilcoxon matched-pairs test. **f, h**, Ratios of the Tfr or Tfh cell NCF with CCL3^+/+^ over CCL3^−/−^ HyHEL10 B cells from the same movie. *, P<0.05, Student’s t-test. **i, j**, Contact duration between Tfr or Tfh and HyHEL10 GC B cells of each genotype undergoing Strict (i) or Non Strict (j) interactions. Red lines represent medians. Data from 5 independent experiments (5 mice) per T cell type. **k, l**, Time histograms for duration of strict (k) and non-strict (l) contacts between Tfr and Tfh cells with WT HyHEL10 GC B cells.

**Supplementary Movie 2**). To take into account the ambiguity of correct identification of B-T cell interactions by 2P imaging, we used both “strict” and “non-strict” definitions of contacts between GC B cells and follicular T cells. By “strict” we define the interactions that based on the cell colocalization analysis in 3D have taken place with high confidence. By “non-strict” interactions we identify all likely interactions identified based on cell proximity, including the “strict” interactions. The data was analyzed in a blinded fashion to avoid possible bias in cell contact definition (**Fig. 6d, Supplementary Movie 3**). We calculated the normalized contact frequency (NCF) of Tfr cells with fluorescent CCL3^+/+^ and CCL3^−/−^ GC B cells by dividing the total number of Tfr cell contacts with CCL3^+/+^ or CCL3^−/−^ GC B cells by the average numbers of fluorescent B cells of each type present in the imaged GCs. Tfr cells’ NCF was lower for CCL3^−/−^ compared to CCL3^+/+^ GC B cells in 5 experiments, independently of the “strict” vs “non-strict” B-Tfr cell contact definition (**Fig. 6e-h**). The differences in Tfr cell contact frequencies with CCL3^+/+^ and CCL3^−/−^ GC B cells were not due to distinct migratory properties of CCL3^+/+^ and CCL3^−/−^ B cells (**Supplementary Fig. 6**). In contrast to Tfr cells, OTII Tfh cell NCF with CCL3^+/+^ vs. CCL3^−/−^ GC B was comparable (**Fig. 6e-h, Supplementary Movie 4**). The ratio of T cells’ NCF with CCL3^+/+^ vs. CCL3^−/−^ GC B cells was significantly lower for Tfh compared to Tfr cells (**Fig. 6f, h)**.

While Tfr cells formed more frequent contacts with CCL3^+/+^ GC B cells, duration of Tfr cell interactions with CCL3^+/+^ and CCL3^−/−^ GC B cells was comparable (**Fig. 6i, j**). Additionally, no significant difference in the duration of Tfh cell contacts with CCL3^+/+^ vs. CCL3^−/−^ GC B cells was observed (**Fig. 6i, j**). The vast majority of Tfr and Tfh cell contacts with GC B cells were shorter than 5 min (**Fig. 6i-l**). However, while a substantial number of both Tfh and Tfr cells also formed more prolonged interactions with GC B cells, no Tfr cells’ interactions with GC B cells exceeding 7.5 min were observed (**Fig. 6k, l, Supplementary Movies 3, 4**).

### Transient depletion of Tregs leads to relative increase in CCL3^+/+^ vs. CCL3^−/−^ HyHEL10 GC B cells at the peak of GC response

To determine whether Tfr cells may act on CCL3-proficient GC B cells and modulate their participation in GC response we sought to determine whether transient depletion of Tregs after formation of GCs could affect relative involvement of CCL3 proficient and deficient B cells in the GC response. In order to test that, we co-transferred CCL3^+/+^ and CCL3^−/−^ HyHEL10 B cells into recipient FoxP3^DTR^ mice, immunized mice to promote HyHEL10 cell entry into GC response and then treated mice with diphtheria toxin (DTx) to promote transient depletion of FoxP3^+^ cells or with PBS for control (**Fig. 7a**). We hypothesized that CCL3 secreted by GC B cells promotes their direct interactions with and inhibition by Tfr cells. In that case, depletion of Tfr cells should lead to increased expansion of CCL3-proficient compared to CCL3-deficient GC B cells (**Fig. 7b**). As expected, treatment of the recipient mice with DTx at 6 d.p.i. led to significant drop in Tfr cells numbers in 3 days (**Fig. 7c, d**) and small increase in the GC B cell numbers (**Fig. 7e**). Consistent with that, upon the transient Tfr cell depletion, we detected a small increase in CCL3^+/+^ HyHEL10 B cell numbers, however CCL3^−/−^ HyHEL10 GC B cells were virtually unchanged (**Fig. 7f**). As a result of this, there was significant enrichment of CCL3^+/+^ vs. CCL3^−/−^ HyHEL10 B cells within GCs (**Fig. 7g**). Therefore, the data is consistent with direct CCL3-dependent inhibition of GC B cells by Tregs at the peak of GC response (**Fig. 7b**).

**Figure 7.**
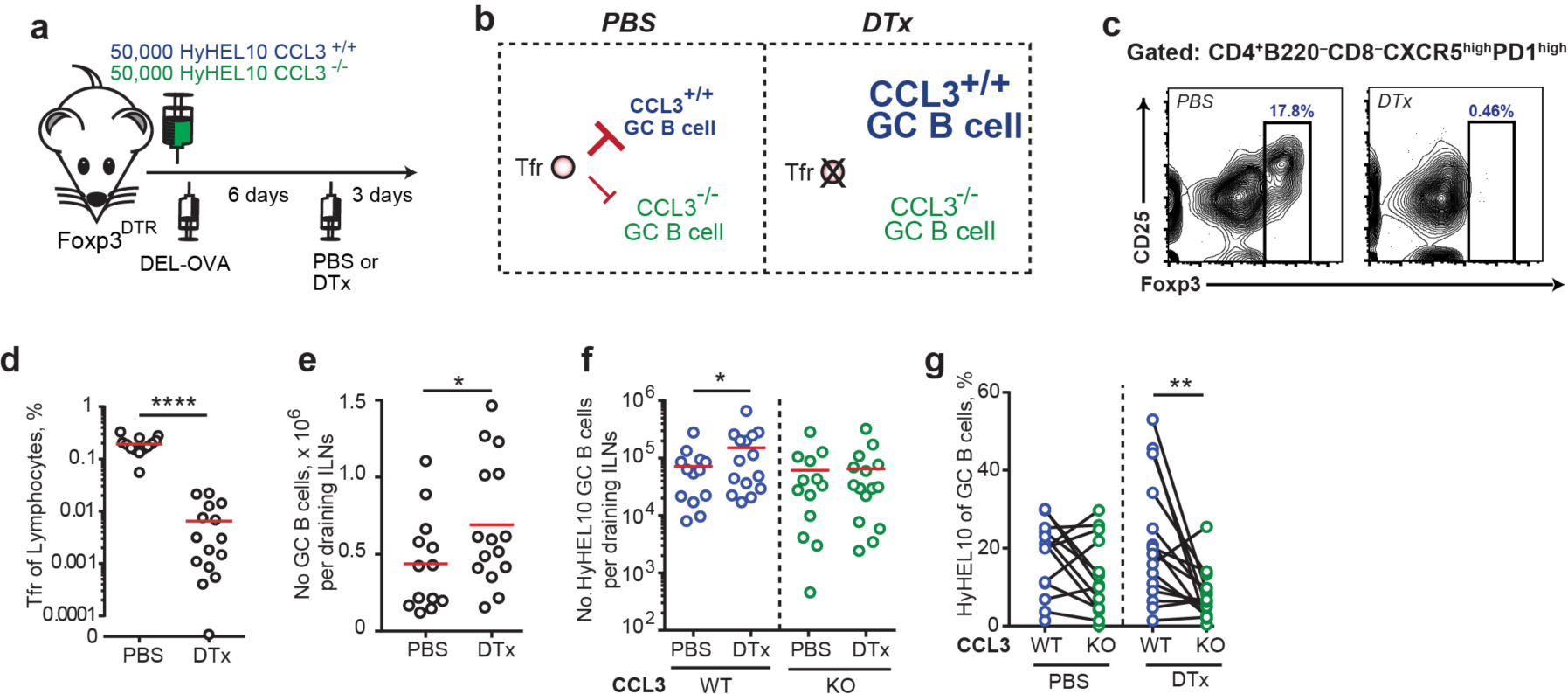
Tregs repress B cell participation in GCs at 9 d.p.i. in a CCL3-dependent way. **a**, Experimental outline. FoxP3^DTR^ recipient mice were transferred with HyHEL10 B cells, immunized with 50 μg DEL-OVA in Ribi s.c. and then treated either with PBS or 5 μg DTx in PBS i.p. 3 days before analysis. **b**, Suggested model of CCL3-dependent regulation of GC B cells by Tfr cells and prediction for CCL3^+/+^ and CCL3^−/−^ GC B cell participation in GCs upon ablation of Tfr cells at 9 d.p.i. **c-g**, Flow cytometry analysis of cell participation in immune response at 9 d.p.i. in DTx-treated and control mice. **c-e**, DTx-mediated depletion of Tfr cells at 9 d.p.i. Representative example of Tfr ablation (in c) and Tfr fraction of total lymphocytes (in d) and increase in GC response in DTx-treated mice (in e). **f, g**, CCL3^+/+^ and CCL3^−/−^ HyHEL10 GC B cells in PBS and DTx-treated mice shown as the total numbers of cells in dLNs (f) or as the % of total GC B cells. The data are from 5 independent experiments. Each dot represents a single mouse and red lines correspond to the mean values. Paired analyses are indicated using solid black line. *, p<0.05, **, p<0.01, ****, P<0.0001. Two-sided (in d, g) or one-sided (in e, f) Student’s t-test.

## DISCUSSION

While Tfr cells have been shown to control the numbers, specificity to foreign antigen and cytokine production of Tfh cells that support GC responses *in vivo* (Sage, Francisco et al. 2013, Sage, Paterson et al. 2014, Wing, Ise et al. 2014, Wu, Chen et al. 2016) whether Tfr cells can directly affect GC B cells *in vivo* has not been definitively demonstrated. While this study does not exclude the role of Tfr cells in the regulation of Tfh cell numbers and the magnitude of foreign-Ag specific response, it suggests that secretion of CCL3/4 by B cells is important for direct Tfr-mediated regulation of GC B cells. First, we determined that GC centrocytes upregulate expression of CCL3/4 compared to centroblasts. We then showed that Tfr cells as well as the other Treg subsets are responsive to CCL3 and CCL4 in transwell assays and demonstrated that CCL3 production by foreign Ag-specific GC B cells increases their sampling efficiency by Tfr cells. Finally, consistent with a model of direct inhibition of CCL3-producing GC B cells by Tfrs, we found that depletion of regulatory T cells at the peak of the GC response results in a small but significant increase in the numbers of CCL3-proficient, but not CCL3-deficient HyHEL10 GC B cells. Future studies should assess which molecular players, including TGF-β, PD1, CTLA-4, GLUT1, and IL21 (Alexander, Tygrett et al. 2011, Sage, Francisco et al. 2013, Sage, Paterson et al. 2014, Wing, Ise et al. 2014, Sage, Ron-Harel et al. 2016, Wu, Chen et al. 2016) may be involved into the observed direct regulation.

Future studies should examine the impact of B cell-intrinsic CCL3 deficiency on the kinetics of B cell participation in the GC and memory response, and foreign Ag-specific Ab responses and dissect the contribution of Tfr cells to the regulation. They should also assess whether CCL3 contributes to better control over bystander or self-reactive B cell clones in GCs and conveys better protection against development of autoimmunity. Interestingly, although CCL3 and CCL4 are coexpressed in activated and GC B cells, neutralization of CCL4 at the time of GC formation did not promote significant increase in the GC response in WT or CCL3-KO mice. It is possible that CCL4 expression in CCL3 KO cells is reduced as has been reported before (Patterson, Pesenacker et al. 2016). Alternatively, CCL4 concentration produced by GC B cells is suboptimal for promotion of Tfr cell response.

We suggest two possible models that could explain the increased sampling of CCL3-proficient GC B cells by Tfr cells. First, the observed effect could be due to local chemoattraction of Tfr cells to CCL3-secreting GC B cells. Since BCR crosslinking has been reported to induce upregulation of CCL3/4 production in GC B cells (Krzysiek, Lefevre et al. 1999), we hypothesize that GC centrocytes that recently acquired antigen from FDCs (and possibly T cell help) are likely to produce more CCL3/4 than other GC B cells, and thus could form local short-range gradients of these chemokines. In that case, to preserve local CCL3 gradient around migrating GC B cell, secreted CCL3 would have to be rapidly internalized and removed by other cells within GCs. Alternatively to the first model, CCL3 chemokine secreted by GC B cells could be transiently immobilized on the surface of selected GC B cells in association with chemokine-binding glycosaminoglycans (Liang, Triandafillou et al. 2016) and may serve to stabilize very transient probing interactions of GC B cells with Tfr cells that are beyond the resolution capabilities of intravital 2P microscopy. Both of these models can lead to decreased efficiency in productive sampling of CCL3 KO B cells by Tfr cells. Future studies should examine whether CCL3 is uniformly upregulated in GC CC or whether there is a subset of cells with elevated expression of CCL3/4. In the later case the advantage that CCL3^hi^ GC B cells may have in contacting Tfr cells may be significantly underestimated.

Future studies should address whether CCR5, CCR1, or both of these receptors are involved in the observed CCL3-mediated regulation and interactions between GC B cells and Tfr cells.

Previous intravital imaging studies of Tfh cell interactions with cognate GC B cells revealed that majority of these encounters are transient (Allen, Okada et al. 2007). They also suggested that a small fraction of the interactions that are more prolonged (> 5-10 min) may be more efficient for productive communication between the cells and for GC B cell selection (Qi, Cannons et al. 2008, Shulman, Gitlin et al. 2014). In this study we found that similarly to Tfh cells, a majority of interactions between foreign-antigen specific GC B cells and natural Tfr cells *in vivo* are shorter than 5 min. Interestingly, Tfr cells also formed a few interactions with GC B cells that were more prolonged. However, while a few Tfh cell contacts with GC B exceeded 7.5 min, none of these have been observed for Tfr cells. This discrepancy may be due to non-cognate interactions or very weak cognate interactions between foreign antigen specific GC B cells and natural Tfr cells. Future studies should directly address whether Tfr cells recognize MHCII/self-peptides on GC B cells via T cell receptors (TCR), and how prevalent these cognate interactions are. In addition, whether cognate interactions of Tfr cells with self antigen-presenting GC B cells exert much stronger negative control of potentially self-reactive GC B cells must be examined, as well as the contribution of CCL3 to that regulation.

In summary, our findings suggest that local CCL3 chemokine production by GC B cells promotes their interactions and direct inhibition by Tfr cells at the peak of GC response.

## MATERIALS AND METHODES

### Mice, immunizations, and bone marrow chimeras

C57BL/6 (B6, WT) mice were purchased from the National Cancer Institute, Charles River or Jackson Laboratories.

B6-CD45.1 (002014), CCL3-KO (002687), β-actin-CFP (004218), UBC-GFP (004353), Stop-tdTomato (007909) and E2a-Cre (003724) mice were from Jackson Laboratories. HyHEL10 (Allen, Okada et al. 2007), MD4 (Goodnow, Crosbie et al. 1988), OTII (Barnden, Allison et al. 1998), Foxp3^EGFP^, and Foxp3^DTR^ mice were from internal colonies. All mice were housed in specific-pathogen free conditions. Relevant mice were interbred to obtain HyHEL10 CFP^+^, HyHEL10 GFP^+^ CCL3-KO, OTII GFP^+^, OTII tdTomato^+^, MD4 CFP^+^, and tdTomato^+^ Foxp3^EGFP^ mice. 6-12 weeks old mice were immunized s.c. with the protein antigens OVA (Sigma), DEL-OVA (produced as previously described (Allen, Okada et al. 2007)), or NP-KLH (Biosearch Technologies), mixed in either Ribi (Sigma) or Complete Freund Adjuvant (CFA, Sigma). In some experiments 50 μg of anti-CCL4 (R&D clone 46907) or isotype control rat Abs (R&D clone 54447) were s.c. administered into the preimmunized mice. All experiments were performed in compliance with federal laws and institutional guidelines as approved by the University Committee on Use and Care of Animals.

### Cell isolation, flow cytometry analysis and cell sorting

Lymphocytes were isolated by homogenizing lymph nodes (LNs) and/or spleens into a single cell suspension in DMEM medium (Corning) containing 2% fetal bovine serum (FBS, Atlanta Biologicals), antibiotics (50 IU/mL of penicillin and 50 *μ*g/mL of streptomycin; Gibco) and 10 mM HEPES (Gibco) and straining through a 70 *μ*m mesh filter (Falcon) in the presence of 20 *μ*g/mL of DNase I (Sigma-Aldrich). Red blood cells were lysed using Tris-buffered NH_4_Cl. The following antibodies and reagents were used for flow cytometry analysis: CD3 (BD, 145-2C11), CD4 (BD, RM4-5), CD8 (BD, 53-6.7), CD25 (BD, PC61.5), B220 (BD, RA3-6B2), CD19 (BD, 1D3), CXCR5 (BD, 2G8), Fas (BD, Jo2), IgM (BD, R6-60.2), IgM^a^ (BD, DS-1), νβ5 (BD, MR9-4), CD43 (BD, S7), CD19 (Biolegend, 6D5), CD45.1 (Biolegend, A20), CD45.2 (Biolegend, 104), IgD (Biolegend, 11-26c.2a), PD-1 (Biolegend, RMP1-30), CXCR4 (eBiosciences, 2B11), CD86 (Biolegend, GL1), Foxp3 (eBiosciences, FJK-16s), GL-7 (eBiosciences, GL-7), SA-qDot607 (Life Technologies), SA-DyLight 488 (Biolegend). Single-cell suspensions were incubated with biotinylated antibodies for 20 minutes on ice, washed twice with 200 *μ*! PBS supplemented with 2% FBS, 1 mM EDTA, and 0.1% NaN (FACS buffer), and then incubated with fluorophore-conjugated antibodies and streptavidin for 20 minutes on ice, and washed twice more with 200 *μ*l FACS buffer. For FoxP3 staining the cells were permeabilized and stained using FoxP3 staining buffer set (eBioscience) according to the manufacturer’s instructions. Cells were then resuspended in FACS buffer for acquisition. All flow cytometry analyses and cell-sorting procedures were done using FACSCanto II and FACSAria IIIu respectively. FlowJo Software (v 9.7; TreeStar) was used for data analyses and plot rendering.

### Cell purification and adoptive transfers

For adoptive transfers, cells were isolated from combined spleens and LNs of donor mice and CD4 T cells or B cells were enriched using autoMACS (Miltenyi Biotec) as described before (Allen, Okada et al. 2007). The purity of B cells was >95%, and CD4 T cells >70% for all experiments. Lymphocytes were adoptively transferred by intravenous injection into the lateral tail vein.

### Generation of mice with Tregs and Tfr cells expressing tdTomato

In order to generate mice with fluorescent Tregs the following scheme was utilized: first, tdTomato expressing mice were crossed with Foxp3^EGFP^ mice. Second, tdTomato^+^Foxp3^EGFP^ Tregs were sorted and adoptively transferred into Foxp3^DTR^ mice where endogenous Tregs were transiently ablated by DTx treatment (Sigma). To sort tdTomato expressing Tregs, the LNs and spleens from the tdTomato^+^Foxp3^EGFP^ mice were combined and lymphocyte suspension was prepared as described above. The lymphocytes were separated from RBCs using Ficoll-Paque (GE Healthcare) gradients per manufacturer’s instructions using 14 mL round bottom tubes (Falcon). Single cell suspensions were enriched for CD4^+^ T cells as described above. Following the enrichment, EGFP^+^ cells were sorted into DMEM medium supplemented with 10% FCS. The purity of sorted Tregs as determined by intracellular Foxp3 staining was > 99%. About 0.8-1.5 million of purified tdTomato^+^ Tregs were then transferred into recipient Foxp3^DTR^ mice via tail vein injection. Finally, one day later the endogenous nonfluorescent Tregs in the recipient Foxp3^DTR^ mice were ablated by intraperitoneal injection of 5 **μ**g/kg of DTx in PBS. The DTx treatment was repeated once more a week later.

### Cell culture and chemotaxis

Transwells with 5 **μ**m pore size (Corning) were used. CD4 T cells were isolated and enriched as described above from draining peripheral LNs of mice s.c. immunized with OVA in Ribi at 10 days following immunization. T cells were resuspended with RPMI 1640 (Corning) supplemented with 2% fatty acid free BSA (Sigma-Aldrich), 10 mM HEPES, 50 IU/mL of penicillin, and 50 mg/mL of streptomycin (HyClone). For chemotaxis analysis the lower chambers of transwells were filled with the same medium mixed with various concentrations of CCL3 or CCL4 chemokines (PeproTech). For chemokinesis analysis both upper and lower chambers of transwells were filled with either 200 ng/mL of CCL3 or with 400 ng/mL CCL4. Transwells with chemokines and resuspended CD4 T cells were incubated at 37°C and 5% CO_2_ for 10 minutes. After that, CD4 T cells were placed in the upper chambers of transwells at 4×10^5^ cells per well and incubated at 37°C and 5% CO_2_ for 3 hours. Two to three replicas per condition have been performed per experiment. The transmigrated fraction of cells was stained and analyzed via flow cytometry. Chemotactic index was calculated as the ratio of cells that transmigrated to chemokine compared to no cytokine control.

### Two Photon Microscopy

Inguinal LNs (ILNs) were either explanted or surgically exposed for intravital imaging and perfused as previously described (Allen, Okada et al. 2007, Grigorova, Schwab et al. 2009). ILNs were imaged with a Leica SP5 II (Leica Microsystems) fitted with a MaiTai Ti:Sapphire laser (Sepctra-Physics) that was tuned to 870 nm. Each *xy* plane spanned 435 **μ**m × 435 **μ**m and with z spacing ranging from 2-3 **μ**m detecting emission wavelengths of 430-450 nm (second harmonic emission of collagen), 465-500 nm (for CFP^+^ cells), 520-550 nm (for GFP^+^ cells), and >560 nm (for tdTomato^+^ cells), every 20-25 seconds. Images were acquired by Leica Advanced Fluorescent Suite (Leica Microsystems). Analysis of the imaging data and generation of 3D rotations and time-lapse image sequences were performed using Imaris 7.6.5 × 64 (Bitplane). Videos were processed with a median noise filter. Semi-automated cell tracking in 3D was performed with Imaris 7.6.5 × 64, and then verified and corrected manually. 3-dimensional GC volume was defined based on the distribution of HyHEL10 CFP B cells by combination of visual analysis and a custom-made MATLAB program that performed time integrated image rendering of CFP signal. TdTomato^+^ Tregs and Th that transited within the follicles and GCs were tracked. Their interaction with WT and CCL3-KO B cells within defined GCs volume were visually identified and categorized either as a strict contact as defined when cell-to-cell contact was unambiguous or a non-strict contacts where cells could be observed in extreme proximity (~1 **μ**m). Finally, we normalized the number of contacts to the average number of WT or CCL3-KO B cells within the GC volume accessible to Tfr or Tfh cells to arrive at a normalized contact frequency. Annotation and final compilation of videos was performed in Adobe After Effects CS5.5 (Adobe).

### RT-PCR Analysis

RNA from sorted cells was obtained using RNeasy Kit (Qiagen) following the manufacturer’s instructions. RNA was treated with DNase to remove genomic DNA (Ambion). The concentration of RNA was calculated using a NanoDrop 2000 (Thermo) and cDNA was synthesized using a SuperScript III kit (Invitrogen) following the manufacturer’s instructions. Preamplification of target genes was performed using PreAmp Kit (AB Biosystems) for 10 cycles. TaqMan assays were obtained from Applied Biosystems and RT-PCR was carried out on a RealPlex 2 (Eppendorf). Expression levels of CCL3/4 were normalized to the level of β2m.

### Statistical analysis

All statistical tests were computed with PRISM (GraphPad) after consultation with a University of Michigan Center for Statistical Consultation and Research representative. Statistical analysis of data normalized to the control samples were performed using a one-sample *t*-test. For comparisons between two groups *t*-test was utilized. Welch’s correction was applied for data with unequal variances. For data in which more than two groups or more than two time points were analyzed, two-way ANOVA followed by Dunnet post-hoc analysis was done. In cases where we did not assume normally distributed data and the data was from paired measurements, we used the Wilcoxon signed-rank test. *P* values of less than 0.05 were considered statistically significant. All statistically significant results were labeled. No samples were excluded from the analysis.

## AUTHOR CONTRIBUTIONS

Z.L.B. planned, performed and analyzed experiments and prepared the manuscript; M.M., R.W. and J.S.T. helped with various aspects of other experiments; J.B.G., M.I.I., and S.S.S. performed blind analysis of the imaging data; I.G. planned, performed and analyzed experiments, and prepared the manuscript.

## ACKNOWLEDGMENTS

We thank W. Dunnick and E. Perkey for discussions; J. Cyster (University of California San Francisco), W. Zhou (University of Michigan Medical school), and A. Rudensky (Memorial Sloan Kettering Cancer Center) for provision of mice; the University of Michigan flow cytometry core for cell sorting assistance. Supported by the American Heart Association (14PRE19960005) to Z.B., by the Herman and Dorothy Miller Award for Innovative Immunology Research to J.S.T. and Z.L.B., and National Institute of Health (R01 AI106806) to I.G. The authors declare no competing financial interests.

## SUPPLEMENTARY FIGURES

**Supplementary Figure 1.**
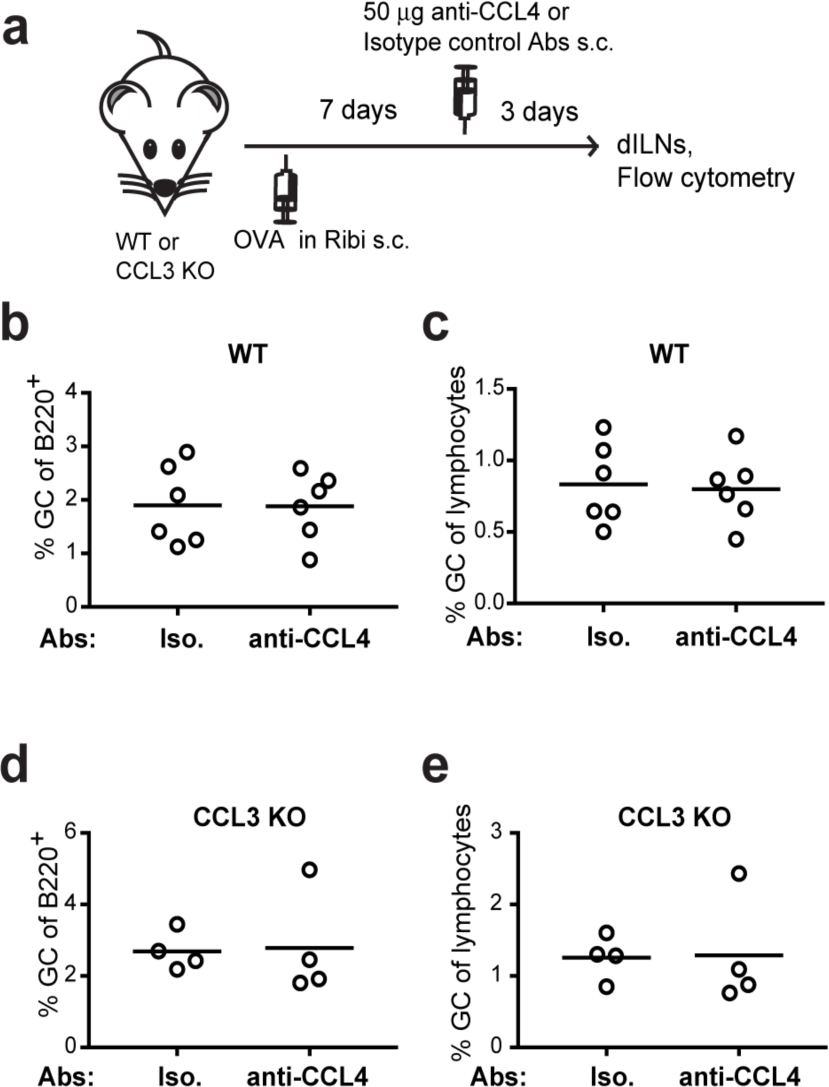
CCL4 neutralization in GCs does not amplify GC response in WT or CCL3-KO mice. a, Experimental strategy, b-e, GC B cells (B220^POS^ CD4^neg^ CD8^neg^ FAS^high^ GL7^high^) in the dLNs of WT (b, c) and CCL3-KO (d, e) mice as a fraction of B220^+^ cells (b, d) or lymphocytes (c, d) at 10 d.p.i. with OVA in Ribi.

**Supplementary Figure 2:**
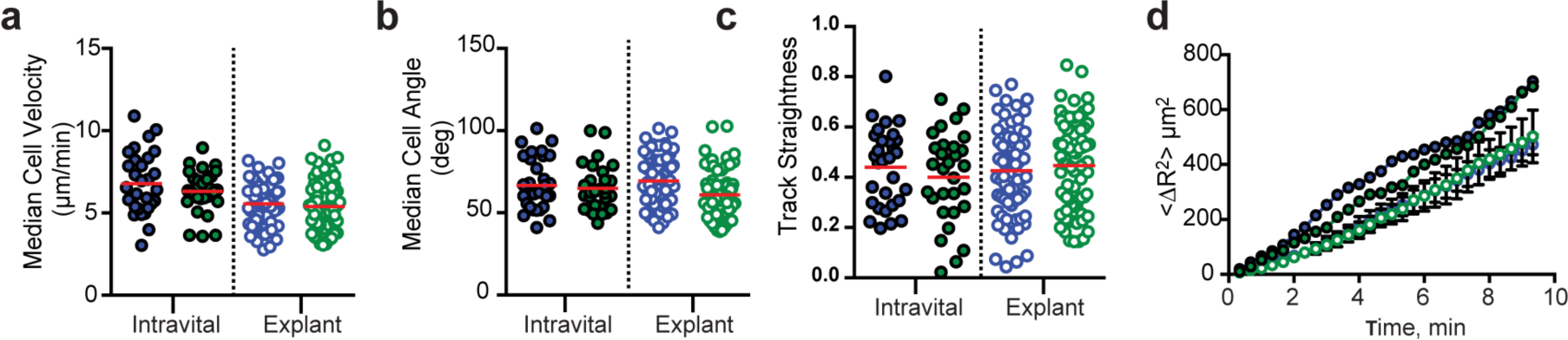
HyHELIO GC CCL3^+/+^ CFP (blue circles) and CCL3^−/−^ GFP (green circles) cell motility analysis. Red lines represent medians. Data are from 4 independent experiments.

## SUPPLEMENTARY MOVIE LEGENDS

**Movie S1. 2-photon imaging of Tfr cells migration in respect to Tfh cells and GC B cells.** Also see **Figure 5e**. A time-lapse sequence of a 110 Mm z-stack of a GC in an explanted inguinal lymph node imaged at 8 days after immunization. HyHEL10 GC B cell; cyan. Tfr; red. Tfh; green. Colored lines indicate the trajectories of the indicated cell types, tracked by Imaris and manually verified. Mice were generated as described in the **Figure 5a**. Time is shown as hh:mm:ss and z-stacks were acquired at 20 second intervals.

**Movie S2. Intravital imaging of Tfr cells in respect to the GCs containing both CCL3^+/+^ and CCL3^−/−^ HyHEL10 B cells.** The data shown corresponds with the images in **Figure 6**. A time-lapse sequence of a GC within inguinal LN of a mouse prepared as described in **Figure 6a** and subjected to intravital imaging at 8 days post immunization. 180 Mm z-stack. GC volume (gray surface) was defined based on the distribution of CFP HyHEL10 cells. Quantitative analysis in **Figure 6e-l** was performed for Tfr cells (red) interactions with HyHEL10 GC B cells (CCL3^+/+^; cyan. cCl3^−/−^; green.) within GC volumes defined in the same fashion. The tracks of individual Tfr cells outside the GC are labeled in purple while interior tracks are represented in yellow lines. Time is expressed as hh:mm:ss and z-stacks were acquired at 20 second intervals.

**Movie S3. Examples of Tfr cells interactions with GC B cells identified as “strict” or “nonstrict” for quantitative analysis.** A time-lapse sequence of a representative Tfr (red) entering the GC (dashed white line) and then undergoing contacts with HyHEL10 CCL3^+/+^ (cyan) or CCL3^−/−^ (green) HyHEL10 GC B cells within inguinal LN of a mouse prepared as described in **Figure 6a** and subjected to 2P imaging at 8 days post immunization. A 40 **μ**Μ slice is in view. Inlets are zoomed in and 3D rotated to visualize the contact. Time is expressed as mm:ss and z-stacks were acquired at 25 second intervals

**Movie S4. 2-photon imaging of Tfh cells interacting with GC B cells**. A time-lapse sequence of a GC within an inguinal LN of a mouse prepared as described in **Figure 6b** and subjected to explant imaging at 8 days post immunization. 140 um z-stack. Quantitative analysis was performed for Tfh cells (red) interactions with HyHEL10 GC B cells (CCL3^+/+^, cyan; CCL3^−/−^, green) within the GC and can be found in **Figure 6e-l**. Examples of long duration (>5 minutes) contacts and short duration (<5 minutes) contacts are shown in 10 um z-projections. Time is shown as hh:mm:ss for both time of the movie and for duration of the indicated contacts. Z-stacks were acquired at 20 second intervals.

## REFERENCES

Alexander, C. M., L. T. Tygrett, A. W. Boyden, K. L. Wolniak, K. L. Legge and T. J. Waldschmidt (2011). “T regulatory cells participate in the control of germinal centre reactions.” Immunology 133(4): 452–468.

Allen, C. D., T. Okada, H. L. Tang and J. G. Cyster (2007). “Imaging of germinal center selection events during affinity maturation.” Science 315(5811): 528–531.

Aloulou, M., E. J. Carr, M. Gador, A. Bignon, R. S. Liblau, N. Fazilleau and M. A. Linterman (2016). “Follicular regulatory T cells can be specific for the immunizing antigen and derive from naive T cells.” Nat Commun 7: 10579.

Barnden, M. J., J. Allison, W. R. Heath and F. R. Carbone (1998). “Defective TCR expression in transgenic mice constructed using cDNA-based alpha- and beta-chain genes under the control of heterologous regulatory elements.” Immunol Cell Biol 76(1): 34–40.

Bettelli, E., Y. Carrier, W. Gao, T. Korn, T. B. Strom, M. Oukka, H. L. Weiner and V. K. Kuchroo (2006). “Reciprocal developmental pathways for the generation of pathogenic effector TH17 and regulatory T cells.” Nature 441(7090): 235–238.

Botta, D., M. J. Fuller, T. T. Marquez-Lago, H. Bachus, J. E. Bradley, A. S. Weinmann, A. J. Zajac, T. D. Randall, F. E. Lund, B. Leon and A. Ballesteros-Tato (2017). “Dynamic regulation of T follicular regulatory cell responses by interleukin 2 during influenza infection.” Nat Immunol 18(11): 1249–1260.

Bystry, R. S., V. Aluvihare, K. A. Welch, M. Kallikourdis and A. G. Betz (2001). “B cells and professional APCs recruit regulatory T cells via CCL4.” Nat Immunol 2(12): 1126–1132.

Caron, G., S. Le Gallou, T. Lamy, K. Tarte and T. Fest (2009). “CXCR4 expression functionally discriminates centroblasts versus centrocytes within human germinal center B cells.” J Immunol 182(12): 7595–7602.

Chung, Y., S. Tanaka, F. Chu, R. I. Nurieva, G. J. Martinez, S. Rawal, Y. H. Wang, H. Lim, J. M. Reynolds, X. H. Zhou, H. M. Fan, Z. M. Liu, S. S. Neelapu and C. Dong (2011). “Follicular regulatory T cells expressing Foxp3 and Bcl-6 suppress germinal center reactions.” Nat Med 17(8): 983–988.

Compagno, M., W. K. Lim, A. Grunn, S. V. Nandula, M. Brahmachary, Q. Shen, F. Bertoni, M. Ponzoni, M. Scandurra, A. Califano, G. Bhagat, A. Chadburn, R. Dalla-Favera and L. Pasqualucci (2009). “Mutations of multiple genes cause deregulation of NF-kappaB in diffuse large B-cell lymphoma.” Nature 459(7247): 717–721.

Cook, D. N., M. A. Beck, T. M. Coffman, S. L. Kirby, J. F. Sheridan, I. B. Pragnell and O. Smithies (1995). “Requirement of MIP-1 alpha for an inflammatory response to viral infection.” Science 269(5230): 1583–1585.

Fu, W., X. Liu, X. Lin, H. Feng, L. Sun, S. Li, H. Chen, H. Tang, L. Lu, W. Jin and C. Dong (2018). “Deficiency in T follicular regulatory cells promotes autoimmunity.” J Exp Med 215(3): 815–825.

Goodnow, C. C., J. Crosbie, S. Adelstein, T. B. Lavoie, S. J. Smith-Gill, R. A. Brink, H. Pritchard-Briscoe, J. S. Wotherspoon, R. H. Loblay, K. Raphael and et al. (1988). “Altered immunoglobulin expression and functional silencing of self-reactive B lymphocytes in transgenic mice.” Nature 334(6184): 676–682.

Grigorova, I. L., S. R. Schwab, T. G. Phan, T. H. Pham, T. Okada and J. G. Cyster (2009). “Cortical sinus probing, S1P1-dependent entry and flow-based capture of egressing T cells.” Nat Immunol 10(1): 58–65.

Kim, J. M., J. P. Rasmussen and A. Y. Rudensky (2007). “Regulatory T cells prevent catastrophic autoimmunity throughout the lifespan of mice.” Nat Immunol 8(2): 191–197.

Krzysiek, R., E. A. Lefevre, W. Zou, A. Foussat, J. Bernard, A. Portier, P. Galanaud and Y. Richard (1999). “Antigen receptor engagement selectively induces macrophage inflammatory protein-1 alpha (MIP-1 alpha) and MIP-1 beta chemokine production in human B cells.” J Immunol 162(8): 4455–4463.

Liang, W. G., C. G. Triandafillou, T. Y. Huang, M. M. Zulueta, S. Banerjee, A. R. Dinner, S. C. Hung and W. J. Tang (2016). “Structural basis for oligomerization and glycosaminoglycan binding of CCL5 and CCL3.” Proc Natl Acad Sci U S A 113(18): 5000–5005.

Lim, H. W., P. Hillsamer, A. H. Banham and C. H. Kim (2005). “Cutting edge: direct suppression of B cells by CD4+ CD25+ regulatory T cells.” J Immunol 175(7): 4180–4183.

Linterman, M. A., W. Pierson, S. K. Lee, A. Kallies, S. Kawamoto, T. F. Rayner, M. Srivastava, D. P. Divekar, L. Beaton, J. J. Hogan, S. Fagarasan, A. Liston, K. G. Smith and C. G. Vinuesa (2011). “Foxp3+ follicular regulatory T cells control the germinal center response.” Nat Med 17(8): 975–982.

Maurer, M. and E. von Stebut (2004). “Macrophage inflammatory protein-1.” Int J Biochem Cell Biol 36(10): 1882–1886.

Menten, P., A. Wuyts and J. Van Damme (2002). “Macrophage inflammatory protein-1.” Cytokine Growth Factor Rev 13(6): 455–481.

Patterson, S. J., A. M. Pesenacker, A. Y. Wang, J. Gillies, M. Mojibian, K. Morishita, R. Tan, T. J. Kieffer, C. B. Verchere, C. Panagiotopoulos and M. K. Levings (2016). “T regulatory cell chemokine production mediates pathogenic T cell attraction and suppression.” J Clin Invest 126(3): 1039–1051.

Qi, H., J. L. Cannons, F. Klauschen, P. L. Schwartzberg and R. N. Germain (2008). “SAP-controlled T-B cell interactions underlie germinal centre formation.” Nature 455(7214): 764–769.

Sage, P. T., L. M. Francisco, C. V. Carman and A. H. Sharpe (2013). “The receptor PD-1 controls follicular regulatory T cells in the lymph nodes and blood.” Nat Immunol 14(2): 152–161.

Sage, P. T., A. M. Paterson, S. B. Lovitch and A. H. Sharpe (2014). “The coinhibitory receptor CTLA-4 controls B cell responses by modulating T follicular helper, T follicular regulatory, and T regulatory cells.” Immunity 41(6): 1026–1039.

Sage, P. T., N. Ron-Harel, V. R. Juneja, D. R. Sen, S. Maleri, W. Sungnak, V. K. Kuchroo, W. N. Haining, N. Chevrier, M. Haigis and A. H. Sharpe (2016). “Suppression by TFR cells leads to durable and selective inhibition of B cell effector function.” Nat Immunol 17(12): 1436–1446.

Shulman, Z., A. D. Gitlin, J. S. Weinstein, B. Lainez, E. Esplugues, R. A. Flavell, J. E. Craft and M. C. Nussenzweig (2014). “Dynamic signaling by T follicular helper cells during germinal center B cell selection.” Science 345(6200): 1058–1062.

Victora, G. D., D. Dominguez-Sola, A. B. Holmes, S. Deroubaix, R. Dalla-Favera and M. C. Nussenzweig (2012). “Identification of human germinal center light and dark zone cells and their relationship to human B-cell lymphomas.” Blood 120(11): 2240–2248.

Wing, J. B., W. Ise, T. Kurosaki and S. Sakaguchi (2014). “Regulatory T cells control antigen-specific expansion of Tfh cell number and humoral immune responses via the coreceptor CTLA-4.” Immunity 41 (6): 1013–1025.

Wing, J. B. and S. Sakaguchi (2014). “Foxp3(+) T(reg) cells in humoral immunity.” Int Immunol 26(2): 61–69.

Wollenberg, I., A. Agua-Doce, A. Hernandez, C. Almeida, V. G. Oliveira, J. Faro and L. Graca (2011). “Regulation of the germinal center reaction by Foxp3+ follicular regulatory T cells.” J Immunol 187(9): 4553–4560.

Wu, H., Y. Chen, H. Liu, L. L. Xu, P. Teuscher, S. Wang, S. Lu and A. L. Dent (2016). “Follicular regulatory T cells repress cytokine production by follicular helper T cells and optimize IgG responses in mice.” Eur J Immunol 46(5): 1152–1161.

